# The plasticity of the pyruvate dehydrogenase complex confers a labile structure that is associated with its catalytic activity

**DOI:** 10.1101/2020.06.02.130369

**Authors:** Jaehyoun Lee, Seunghee Oh, Saikat Bhattacharya, Ying Zhang, Laurence Florens, Michael P. Washburn, Jerry L. Workman

**Affiliations:** Stowers Institute for Medical Research, Kansas City, MO 64110, USA; Department of Pathology and Laboratory Medicine, University of Kansas Medical Center, Kansas City, KS 66160, USA

**Keywords:** PDC, mitochondrial metabolism, TCA cycle, pyruvate, acetyl-CoA, high-performance liquid chromatography (HPLC), size exclusion chromatography (SEC)

## Abstract

The pyruvate dehydrogenase complex (PDC) is a multienzyme complex that plays a key role in energy metabolism by converting pyruvate to acetyl-CoA. An increase of nuclear PDC has been shown to be correlated with an increase of histone acetylation that requires acetyl-CoA. PDC has been reported to form a ~ 10 MDa macromolecular machine that is proficient in performing sequential catalytic reactions via its three components. In this study, we show that the PDC displays size versatility in an ionic strength-dependent manner using size exclusion chromatography of yeast cell extracts. Biochemical analysis in combination with mass spectrometry indicates that yeast PDC (yPDC) is a salt-labile complex that dissociates into sub-megadalton individual components even under physiological ionic strength. Interestingly, we find that each oligomeric component of yPDC displays a larger size than previously believed. In addition, we show that the mammalian PDC also displays this uncommon characteristic of salt-lability, although it has a somewhat different profile compared to yeast. We show that the activity of yPDC is reduced in higher ionic strength. Our results indicate that the structure of PDC may not always maintain its ~ 10 MDa organization, but is rather variable. We propose that the flexible nature of PDC may allow modulation of its activity.

## 1. INTRODUCTION

The multienzyme pyruvate dehydrogenase complex (PDC) catalyzes the reaction that generates acetyl-CoA from pyruvate, the end-product of glucose breakdown. As such, the PDC plays a central role as a gatekeeper of energy and glucose homeostasis. Consistent with its important role in metabolism, low PDC activity arising from mutations of its components causes a PDC deficiency (PDCD) [1]. PDCD patients have a low survival rate and suffer from ataxia and neurodevelopmental delay. This arises from a failure to produce enough energy as the tricarboxylic acid (TCA) cycle, which links glycolysis to electron transport chain for ATP synthesis, needs PDC to convert pyruvate into acetyl-CoA [1,2]. Being a key enzyme in cellular metabolism, PDC is also considered a target for anticancer and antibacterial drugs [3,4]. Furthermore, the level of nuclear PDC is correlated with that of histone acetylation as well as its recruitment to some promoters upon induction possibly for local acetyl-CoA production [5–8].

The central role of PDC in energy homeostasis necessitates a tight regulation of its activity. Short-term regulation (minutes to hours) occurs through the inhibitory phosphorylation of E1α by pyruvate dehydrogenase kinases (PDKs) and phosphatases, while long-term regulation of PDC activity can be exerted at transcriptional levels within days to weeks [9,10]. PDKs have been studied as potential drug targets as well because of their upregulation or activation in diseases such as diabetes and cancer [11,12].

The catalysis of PDC is performed by three separate enzymes E1p (pyruvate decarboxylase), E2p (dihydrolipoamide acyltransferase), and E3 (dihydrolipoamide dehydrogenase), which are linked together efficiently into a large multienzyme complex, where the oligomeric E2p forms a structural core, to which multiple copies of the E1p, E3, and E3BP (E3 binding protein) are bound [13]. The E1p performs the rate-limiting oxidative decarboxylation of pyruvate via thiamine pyrophosphate (TPP) and transfers the acetyl-group to the lipoyl group of the E2p. Then the E2p transfers the acyl group to CoA while reducing its lipoyl domain, which is then oxidized by the E3 which transfers electrons to NAD^+^ via its cofactor FAD, resulting in the formation of NADH [14]. The formation of PDC not only allows substrate channeling between neighboring enzymes but also connects remote E1p and E3 within the complex by active-site coupling via flexible lipoyl domains of the E2p core [15–17].

Attempts to study the PDC has primarily used structure analysis methods such as analytical ultracentrifugation, Cryo-EM, crystallography, isothermal titration calorimetry, nuclear magnetic resonance spectroscopy, small-angle X-ray scattering and neutron scattering [18–24]. Interestingly, structural studies of the PDC have reported varying sizes of the complex [25–30]. In this study, we used size exclusion chromatography (SEC) to examine the size of PDC in its native condition. SEC is a well-established flow-assisted separation technique that can determine protein heterogeneity by separating native protein complexes in solution assisted by mass spectrometry [31]. Although SEC does not provide a detailed molecular information, it can determine hydrodynamic sizes of macromolecules from which oligomerization state(s) may be deduced. For a globular oligomeric complex like the PDC that may organize into multiform structures, SEC can give a good approximation of their sizes.

In this work, by using calibrated SEC using a gel filtration column, we demonstrate that the PDC displays size versatility in an ionic strength-dependent manner. Also, by performing immunoprecipitation followed by mass spectrometry along with SEC, we show that the individual PDC components can form sub-megadalton complexes in a physiological condition. We examine the biological implications of these findings by showing that the activity of PDC is reduced in high ionic strength. We propose that the versatile nature of PDC structure may allow modulation of its activity and translocation.

## 2. Materials and Methods

### 2.1. Yeast strains and growth medium

*Saccharomyces cerevisiae* strains (BY4741) used in this study are listed in Table S1. All single deletion mutants using KanMX4 marker were obtained from Open Biosystems library maintained by Stowers Institute Molecular Biology facility. The remaining strains were generated by targeted homologous recombination of PCR fragments with marker genes. The deletion or tagged strains were verified by PCR with primer sets specific for their deletion or tagging marker or coding regions. Growth and maintenance of *S. cerevisiae* strains were done in Yeast extract–Peptone– Dextrose medium (YPD, 2% glucose).

### 2.2. Yeast cell extractions and subcellular fractionation

Yeast cells were grown in YPD medium at 30°C to OD600 of 0.8. Cell pellet was washed and resuspended in 40 ml per 1,000 OD600 of SB buffer (1.4 M Sorbitol, 40 mM HEPES, pH 7.5, 0.5 mM MgCl_2_, 10 mM β-mercaptoethanol) and spheroplasted with 2 mg/ml Zymolyase 20T (Amsbio LLC, 120491-1) at 23 °C for 45 min. For whole cell extract, spheroplast was lysed with glass beads in TAP50 buffer (50 mM NaCl, 40 mM HEPES pH 7.5, 10 % Glycerol, 0.1 % Tween-20, 1 μg/ml pepstatin A, 2 μg/ml leupeptin, 1 mM PMSF) or EM150 (150 mM NaCl, 10 mM MOPS buffer pH 7.2, 1 mM EDTA, 1 % Triton X-100, 1 μg/ml pepstatin A, 2 μg/ml leupeptin, 1 mM PMSF) at 4 °C for 20 min. The whole cell extract was treated with 25 unit of Benzonase (Millipore Sigma, 70664) and 5 μg of heparin per 1,000 OD cells at 23 °C for 15 min to remove nucleic acid contamination and then clarified by ultracentrifugation at 45,000 rpm for 90 min following a centrifugation at 20,000 g for 20 min. For the separation of high sedimentation rate fraction (HS) and low sedimentation rate fraction (LS) fractions, spheroplast was resuspended in 20 ml of homogenization buffer (10 mM Tris-HCl pH 7.4, 0.6 M sorbitol, 1 mM EDTA, 0.2 % (w/v) BSA) and homogenized with 8 strokes of a revolving pestle at full speed per 1,000 OD600 cells. The homogenized spheroplast was centrifuged at 1,500 g for 5 min and the resulting pellet was collected as the LS fraction. The supernatant was further centrifuged at 3,000 g for 5 min then at 12,000 g for 15 min, when the final pellet was collected as the HS fraction. The HS fraction was washed with 6 ml of buffer C (0.25 M sucrose, 10 mM Tris pH 6.7, 0.15 mM MgCl_2_) and D (0.25 M sucrose, 10 mM Tris pH 7.6, 10 mM EDTA), then finally resuspended in 1 ml of EM buffer (10 mM MOPS buffer pH 7.2, 1 mM EDTA, 1 % Triton X-100, 1 μg/ml pepstatin A, 2 μg/ml leupeptin, 1 mM PMSF) with 0 mM (EM), 150 mM (EM150), or 350 mM NaCl (EM350) and homogenized with 50 strokes and incubated for 1 hr in ice with 5 units of Benzonase. The crude HS fraction was clarified by ultracentrifugation at 50,000 rpm for 30 min. The LS fraction was further clarified by washing the pellet with NP buffer (0.34 M sucrose, 20 mM Tris pH 7.5, 50 mM KCl, 5 mM MgCl_2_) and homogenizing in H350 buffer (40 mM HEPES-KOH pH7.5, 10 % glycerol, 350 mM KCl, 2 mM MgCl_2_, 1 mM EDTA, 0.02% NP40), EM, or EM150 buffer with 25 strokes. The crude extract was incubated with 125 units of Benzonase on a roller for 1 hr in 4 °C, then clarified by ultracentrifugation at 45,000 rpm for 90 min following a centrifugation at 20,000 g for 15 min.

### 2.3. FLAG purification

Clarified extract was incubated with anti-FLAG M2 agarose (Sigma-Aldrich, A2220) at 4 °C for 4 hr and washed two times with its resuspending buffer and once with E100 buffer (25 mM HEPES-KOH pH7.5, 10 % glycerol, 100 mM KCl, 2 mM MgCl2, 1 mM EDTA with Roche cOmplete™, EDTA-free Protease Inhibitor Cocktail Tablets). The FLAG tagged proteins were eluted with E100 buffer containing 400 μg/ml 3xFLAG peptide.

### 2.4. Mammalian protein extraction and Halo purification

For Halo (HaloTag, Promega) purification, HEK293T cells at 40 % confluency in a 100 mm plate were transfected with 10 μg of plasmid containing ORF of PDHA1-HA-Halo or Halo-PDHA1 in 1X BBS (from 2XBBS (0.28 M NaCl, 0.05 M N,N-bis-(2-Hydroxyethyl)-2-aminoethanesulfonic acid,1.5 mM Na_2_HPO_4_)) with 125 mM CaCl_2_. Cells were harvested by scraping in PBS following a PBS wash after 24 hr from transfection. For non-transfected cells, cells were harvested at 100 % confluency. Cell pellet was resuspended in 1 ml of Hypotonic buffer (50 mM Tris-HCl pH 7.5, 0.1 % NP-40) and centrifuged at 4000 rpm for 5 min. The resulting pellet was resuspended in 1 ml of lysis buffer (50 mM Tris-HCl pH 7.5, 1 % Triton X-100, 0.1 % Na-deoxycholate) with 0 mM, 150 mM, or 350 mM NaCl then clarified by centrifugation at 12,000 rpm for 30 min. The crude extract in the supernatant was treated with 125 units of benzonase for 20 min in 4 °C, then clarified by centrifugation at 12,000 rpm for 10 min. The clarified extract was either directly loaded onto Superose 6 column following pre-clearing with 150 μl of Sepharose CL-2B (GE Healthcare, 17-0140-01) or subjected to Halo purification by incubating with 100 μl of HaloLink™ Resin (Promega, G1915) suspension overnight in 4 °C on a rotator. The beads were washed 3 times then subjected for protein elution with 50 units of AcTEV protease (Invitrogen, 12575-015) in PBS buffer by 8 hr incubation in 4 °C on a rotator.

### 2.5. Size-exclusion chromatography (SEC)

The size exclusion fractionation using Superose^®^ 6 10/300 GL (GE17-5172-01) was performed as previously described [32] with minor modification. Cell extracts followng a pre-clearing with 150 μl of Sepharose CL-2B or elution from FLAG of Halo purification was loaded onto a Superose 6 size-exclusion column (Amersham Bioscience) equilibrated with EM, EM50, or EM150 buffer for 0 mM, 50 mM, or 150 mM NaCl condition for yeast samples, PBS buffer or PBS minus NaCl and KCl for 150 mM or 0mM NaCl mammalian samples, or superose 6 buffer (350 mM NaCl, 40 mM HEPES pH 7.5, 5 % glycerol, 0.1 % Tween 20) for 350 mM NaCl condition. The 500 μl fractions eluted from the column were analyzed by western blots and multidimensional protein identification technology as described in the results section.

### 2.6. Western blots

SDS samples were run on 10 % SDS-PAGE and transferred to 0.45 μm Immobilon-P PVDF Membrane (Millipore Sigma, IPVH00010) and immunoblotted using anti-FLAG (Sigma-Aldrich, A8592) at 1:10,000 dilution in 5 % non-fat milk, anti-HA (Thermofisher Scientific, 26183-HRP) at 1:2,000 dilutions in 5 % non-fat milk, anti-V5 (Abcam, ab9116) and anti-MTCO2 (Abcam, ab110271) at 1:1,000 dilutions in 5% non-fat milk, anti-Lamin A (Abcam, ab26300) and anti-E1α (Abcam, ab110330) at 1:500 dilution in 5 % BSA (Sigma-Aldrich, a9647), anti-E1β (Abcam, ab155996), anti-E2 (Santa Cruz Biotechnology, sc-271534), anti-E3 (Santa Cruz Biotechnology, sc-365977), anti-BAF155 (Santa Cruz Biotechnology, sc-9746) at 1:2,000 dilution in 5% BSA, or anti-Brg1 (Abcam, ab110641) at 1:2,000 dilution in 5% BSA.

### 2.7. PDC activity assay

Iodonitrotetrazolium chloride (INT)-based colorimetric assay was performed base on previous publication [33] with minor modification. Briefly, yPDC was purified with FLAG M2 agarose from whole cell extract of *pkp2*Δ/Pdb1-5xFLAG as described above. From 250 μl elution of FLAG purification of Pdb1, 6 μl was used for the assay. Assay was performed in 20 μl reaction with 50 mM potassium phosphate buffer pH 7.5, 2.5 mM β-Nicotinamide adenine dinucleotide (NAD^+^; Sigma Aldrich, N8410), 5 mM sodium pyruvate (Gibco, 11360070), 0.2 mM Thiamine pyrophosphate (TPP; Sigma Aldrich, C8754), 0.6 mM or 0.4 mM Iodonitrotetrazolium chloride (INT) (Sigma-Aldrich, I8377), 6.5 μM Phenazine ethosulfate (PES; Sigma-Aldrich, P4544), 0.1 mM Coenzyme A trihydrate (VWR, IC10480950), 1 mg/ml BSA (Sigma-Aldrich, A9647), and 1 mM MgCl_2_ with 50 mM, 100 mM or 350 mM NaCl. The amount of INT-formazan reduced by PES which was reduced by NADH from PDC activity was measured by the absorbance value at 490 nm by NanoDrop 2000 (Thermofisher, ND-2000).

### 2.8. Proteomics analysis by Multidimensional Protein Identification Technology (MudPIT)

TCA-precipitated protein pellets were solubilized using Tris-HCl pH 8.5 and 8 M urea, followed by addition of TCEP (Tris(2-carboxyethyl)phosphine hydrochloride; Thermo Scientific, 20490) and chloroacetamide (CAM; Sigma-Aldrich, 22790) to final concentrations of 5 mM and 10 mM, respectively. Proteins were digested using Endoproteinase Lys-C at 1:100 w/w (Roche, 11058533103) at 37 °C overnight. Digestion solutions were brought to a final concentration of 2 M urea and 2 mM CaCl_2_ and a second digestion was performed overnight at 37 °C using trypsin (Promega, v5280) at 1:100 w/w. The reactions were stopped using formic acid (5 % final). Peptide samples were loaded on a split-triple-phase fused-silica micro-capillary column and placed in-line with a linear ion trap mass spectrometer (LTQ) (Thermo Scientific), coupled with a Quaternary Agilent 1100 Series HPLC system. A fully automated 10-step chromatography run (for a total of 20 hr) was carried out, as described in [34]. Each full MS scan (400–1600 m/z) was followed by five data-dependent MS/MS scans. The number of the micro scans was set to 1 both for MS and MS/MS. The dynamic exclusion settings used were as follows: repeat count 2; repeat duration 30 sec; exclusion list size 500 and exclusion duration 120 sec, while the minimum signal threshold was set to 100.

MS/MS peak files were extracted with RawDistiller [35] and searched using ProLuCID (v. 1.3.3) [36] against a database consisting of 5945 S. cerevisiae non-redundant proteins (NCBI, 2017-05-16), 193 usual contaminants (such as human keratins, IgGs, and proteolytic enzymes), and, to estimate false discovery rates (FDRs), 6138 randomized amino acid sequences derived from each non-redundant protein entry. The human dataset was searched against a database consisting of 36,628 Homo sapiens NR proteins (NCBI-released June 10, 2016), 193 usual contaminants, and randomized amino acid sequences derived from each NR protein entry. Mass tolerance for precursor and fragment ions was set at 800 ppm. ProLuCID searches were set up against a preprocessed database of tryptic peptides with K/R at both ends and with a differential modification of+16 Da on methionine residues. No maximum number of missed cleavages were specified. To account for alkylation by CAM, 57 Da were added statically to the cysteine residues.

Peptide/spectrum matches were sorted and selected using DTASelect/CONTRAST (v. 1.9) [37] in combination with an in-house software, swallow (v. 0.0.1, https://github.com/tzw-wen/kite), to filter spectra, peptides, and proteins at FDRs < 5 %. Combining all runs, proteins had to be detected by at least 2 peptides or by 1 peptide with 2 independent spectra. To estimate relative protein levels, distributed normalized spectral abundance factors (dNSAF) were calculated for each detected protein, as described in [38]. The MS datasets for the human and yeast analyses may be obtained from the MassIVE repository with accessions MSV000085125 and MSV000085126, respectively (with password: JLE01146) and from the ProteomeXChange with accessions PXD018125 and PXD018126.

## 3. RESULTS

### 3.1. Yeast PDC components display salt-lability

We analyzed the size of the yeast PDC by size exclusion chromatography (SEC). To facilitate the detection of the PDC components, yeast BY4741 strains were generated in which combinations of the E1p β subunit (E1pβ), E2p and E3 were epitope-tagged at their endogenous locus with 3xHA, V5 and 5xFLAG, respectively. Next, whole-cell extracts of these strains were prepared and loaded onto a Superose 6 column for SEC. The SEC fractions were analyzed by immunoblotting using antibodies against the epitope tags on the PDC components (Fig. S1a). Interestingly, all the three PDC components eluted at two peak fractions belonging to different sizes; one in the void volume at 8 ml (≥ 2 MDa) as expected for its well-known macromolecular structure and the other in the range of 14-15 ml elution volume. The two clear peak elution volumes suggested a heterogeneity in the size of the PDC components, hence we decided to investigate this further.

Recent reports have demonstrated the existence of PDC in the nucleus of mammalian cells which is a non-canonical location for this mitochondrial complex [5–7]. We were curious whether the two elution peaks observed resulted from PDC that belonged to the two different subcellular organelles. To address this, we first tried to fractionate subcellular organelles by differential sedimentation rates upon centrifugation [39]. Indeed, high sedimentation rate fraction (HS) was enriched with mitochondria, while low sedimentation rate fraction (LS) was enriched with nuclei (Fig. S1b). Each of the two enriched extracts was size-fractionated by SEC, and the fractions were collected and probed for PDC components by immunoblotting against their epitope tags (Fig. 1a-b). From the (mitochondria-enriched) HS extract, the elution of the PDC components peaked at 8 ml for several MDa (≥ 2 MDa) as well as at 11-12 ml for ~1 MDa proteins (Fig. 1a), while from the (nuclei-enriched) LS fraction, the PDC components were detected at 14-15 ml elution volume for ~230-440 kDa sized proteins (Fig. 1b).

**Figure 1.**
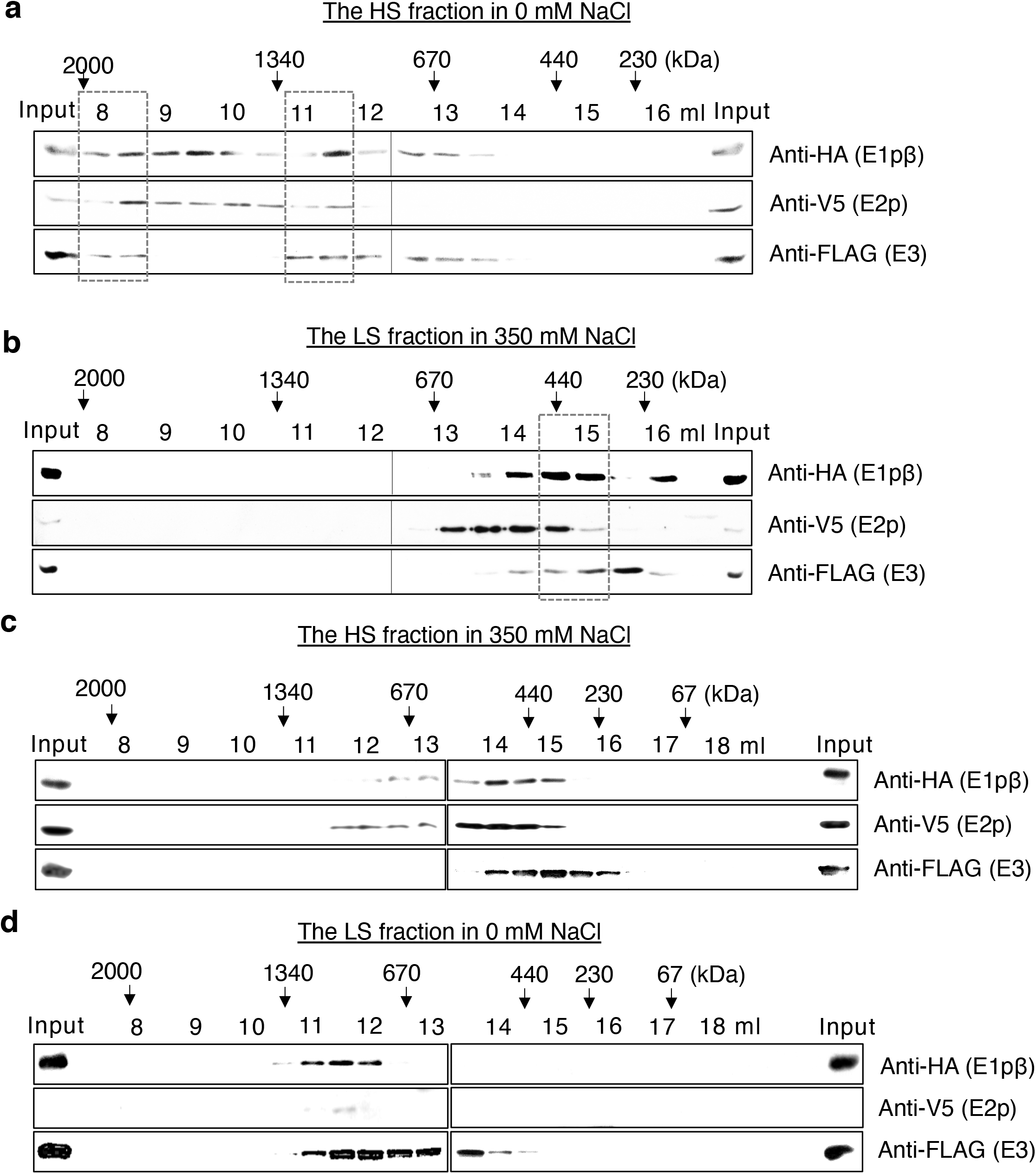
Ionic strength dictates the elution profiles of yPDC components in size exclusion chromatography. Size fraction profiles of yPDC components of Pdb1-3xHA/Lat1-V5/Lpd1-5xFLAG strain from (a) the HS fraction in a buffer containing 0 mM NaCl, (b) the LS fraction in a buffer containing 350 mM NaCl, (c) the HS fraction in a buffer containing 350 mM NaCl, and (d) the LS fraction in a buffer containing 0 mM NaCl. Elution volumes in the SEC via Superose 6 column for every other 500 μl fraction are indicated with the expected size of eluted proteins based on standard proteins. Boxes in dashed lines highlight the peak fractions.

These results seemed to suggest that the size of the PDC differed when extracted from the different subcellular organelles at a first glance, however, the difference in ionic strength between the buffers for preparing the fractions (0 mM for HS fractions vs 350 mM for LS) may have affected the apparent size difference observed. To address this possibility, HS and LS extracts were prepared in the inversed salt conditions. Consequently, HS extract was prepared in the presence of 350 mM NaCl, whereas the LS extract was prepared in 0 mM NaCl. Then each extract was loaded onto a sizing column in the corresponding ionic strength. The eluted fractions from SEC were collected and the epitope-tagged PDC components were probed by immunoblotting (Fig. 1c-d). All the PDC components of HS extract in 350 mM NaCl mostly eluted at 14-15 ml corresponding to molecular weight of ~230-440 kDa (Fig. 1c), whereas those of LS extract in 0 mM NaCl eluted peak at 11-12 ml elution volume for ~1 MDa (Fig. 1d). This result suggested that indeed the PDC elution profile of HS fraction acquired a similar profile to that of LS fraction in the same NaCl concentration in SEC. These data suggested that the size of the PDC components can vary depending on ionic strength, rather than their source of origination.

So far, we have observed that the elution of PDC components in low ionic strength (0 or 50 mM NaCl) appears very different from that in high ionic strength (350 mM NaCl) on a sizing column. Next, we inquired how the elution profile would behave in a physiological salt concentration of 150 mM NaCl. For this, whole-cell extract was prepared with a buffer containing 150 mM NaCl and subjected to SEC. The size-fractions were collected and immunoblotted against each PDC component via its epitope tag (Fig. S1c). Interestingly, all the PDC components in this condition peaked at the elution volume of 14-15 ml for ~230-440 kDa. This was clearly shifted to a smaller sized fraction in 150 mM NaCl compared to the peak elution in the low ionic strength, at 8 ml in 0 mM and 50 mM NaCl, and at 11-12 ml elution volume in 0 mM NaCl (Fig. 1a), suggesting the decreased size of the PDC components in the physiological salt condition. Notably, the size of the small PDC components was reminiscent of that observed in the higher ionic strength (350 mM NaCl) (Fig. 1b-c). Further, MDa sized PDC components that were observed in lower ionic strength (Fig. 1a, S1a) were substantially decreased in the ionic strength of 150 mM NaCl.

Next, to verify the ionic strength-dependent size of the PDC components irrespective of the origin of the extract, HS and LS extracts were prepared and subjected to SEC in 150 mM NaCl. The size-fractions were collected and probed for the PDC components by immunoblotting against their epitope tags (Fig. S1d-e). As in the SEC of whole-cell extract (Fig. S1c), the elution peaks of the PDC components in both HS and LS extracts were shifted to a smaller size of 14-15 ml elution volume comparable to that in the high ionic strength (350 mM NaCl) buffer (Fig. 1b-c). These results indicate that the molecular weight of the PDC components responds to changes in the ionic strength of buffer. In fact, a physiological ionic strength (150 mM NaCl) is enough to form sub-megadalton PDC components.

### 3.2. PDC is a salt labile complex

Thus far, we observed that the size of the PDC components is dependent on ionic strength using SEC of various extracts (Fig. 1 and Fig. S1). To examine whether the purified complex also exhibits a similar behavior, we decided to affinity-purify PDC. For this, strains each expressing a FLAG-tagged E1pβ and E3, were generated. Next, the PDC components were affinity-purified (AP) from cell extracts and the purified complexes were subjected to a mass spectrometry technique known as multidimensional protein identification technology (MudPIT) [34]. Consistent with the fact that E1p, E2p, and E3 interact with one another in the complex, MudPIT revealed that all the PDC components were detected in each of the purifications (Fig. S2a). Next, the purified complexes via E1pβ-5xFLAG (Fig. 2a) and E3-5xFLAG (Fig. 2b) were size-fractionated in 350 mM NaCl and the fractions were probed for their elution profile by immunoblotting against FLAG. Indeed, both purifications of PDC via E1pβ and E3 showed a very similar elution profile to what we observed from the extracts in the same ionic strength (Fig. 1b-c). This result suggested that the purified PDC is as salt-labile as its components.

**Figure 2.**
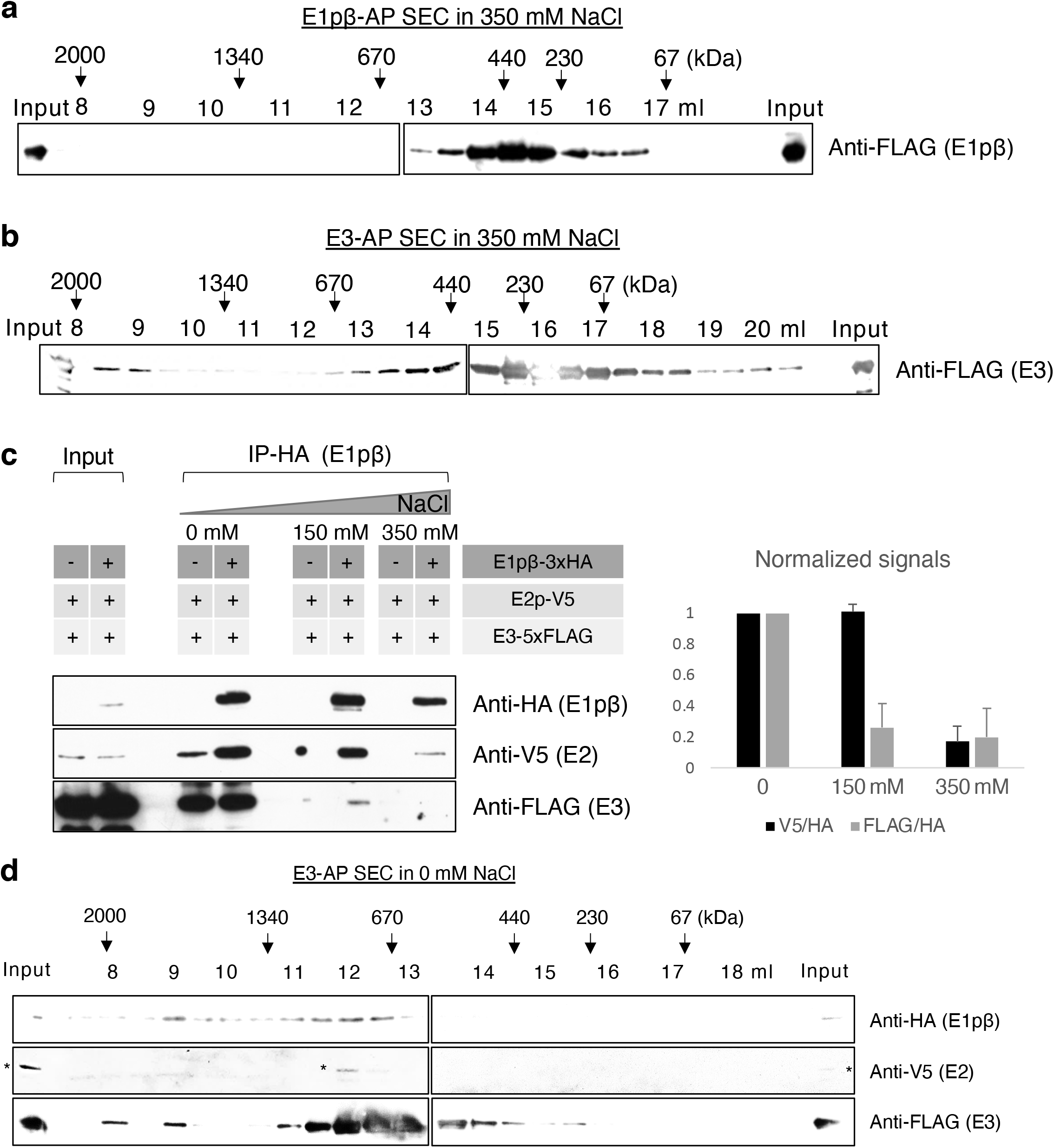
Affinity-purified PDC in SEC and IP suggests that yPDC is a salt-labile complex. (a) Size fraction profiles of FLAG affinity purified (AP) yPDC-E1β from Pdb1-5xFLAG strain in a buffer containing 350 mM NaCl. (b) Size fraction profiles of FLAG purified yPDC-E3 from Lpd1-5xFLAG strain in a buffer containing 350 mM NaCl. (c) (Left panel) Immunoprecipitation (IP) of yPDC-E1β via HA from Pdb1-3xHA/Lat1-V5/Lpd1-5xFLAG strain in a buffer containing 0 mM, 150 mM, or 350 mM of NaCl in EM buffer (10 mM MOPS pH7.2, 1mM EDTA, 1 % Triton-X100). (Right panel) The relative ratios of anti-V5 and anti-FLAG to anti-HA signals are normalized to that of 0 mM and presented with error bars indicating standard deviation of three biological replicates. (d) Size fraction profiles of FLAG AP yPDC-E3 from Pdb1-3xHA/Lat1-V5/Lpd1-5xFLAG strain in a buffer containing 0 mM NaCl. Asterisks denote non-specific signals. For (a), (b), and (d) elution volumes in the SEC via Superose 6 column for every other 500 μl fraction are indicated with the expected size of eluted proteins based on standard proteins.

Since the canonical PDC contains multiple copies of E1p, E2p, and E3 subunits, we set out to determine whether the interactions among the subunits were reduced with increasing ionic strength. If so, this result would be consistent with the decreased size of the complex observed. For that, E1pβ was immunoprecipitated (IP) via HA from whole-cell extract in varying ionic strengths and the amount of the E2p and E3 subunits that co-precipitated were examined by immunoblotting for their epitope tags (Fig. 2c). Indeed, the amount of co-precipitated E2p and E3 decreased with increasing ionic strength, consistent with the decreased size observed by SEC. This result suggests that the PDC in higher ionic strength might contain fewer copies of one or more subunits and thereby, have a smaller size.

To substantiate our findings about the ionic strength-dependent size of PDC further, we set out to detect the dissociation of PDC with increasing ionic strength. For this, we immunoprecipitated PDC via E1pβ-3xHA from whole-cell extract in 0 mM NaCl and performed sequential salt washing on-bead with increasing NaCl concentration from 20 mM to 2 M. To differentiate non-specific binding, a control, E2p-V5, that did not contain HA-tagged E1pβ was also subjected to HA-immunoprecipitation and sequential salt washing. The salt-extracted fractions were collected and immunoblotted against HA and V5 for PDC-E1pβ and E2p, respectively, to monitor the dissociated PDC by the sequential salt-washing (Fig. S2b). Up to 100 mM NaCl condition, non-specifically bound E2p was detected in the salt-extracted fractions, as shown in the V5 immunoblot of both control and E1pβ-HA tagged strains. From the 150 mM NaCl condition, salt-extracted E2p was detected more from the specifically bound PDC via E1pβ-3xHA than from the non-specifically bound E2p from the control. These results suggested that once the concentration of NaCl reached 150 mM, the PDC started dissociating. This was consistent with our SEC results where the main size of the PDC shifted to sub-megadalton in the physiological condition of 150 mM NaCl (Fig. S1c-e), whereas MDa sized PDC was observed in 0 mM (Fig. 1a,d) and 50 mM NaCl (Fig. S1a). The HA-bound PDC-E1pβ was also found in salt-extracted fractions with similar pattern as E2p, further supporting the salt-lability of the PDC (Fig. S2b). Collectively, these results suggest that the PDC is a salt-labile complex which can dissociate in physiological salt conditions (150 mM NaCl). This is in contrast to many protein complexes (e.g. SAGA, SWI/SNF, nucleosome, etc) that are stable in similar conditions [40].

To further validate whether the different sizes of the PDC components observed from cell extracts resulted from the varying size of the complex, we purified the PDC via E3-5xFLAG followed by SEC in 0 mM NaCl condition. The presence of the complex was detected by immunoblotting against the epitopes of the co-purified PDC components (Fig. 2d). Indeed, the elution profiles of the purified PDC matched well with those obtained from extracts (Fig. 1a), suggesting that the complex was eluted with a higher molecular weight (≥ 2 MDa and ~1 MDa) in the lower ionic strength condition. Interestingly, while the E3-AP followed by SEC revealed the complex of MDa sizes (Fig. 2d), it was missing the elution peak of ~230 kDa at 15-16 ml elution volume that was present in the SEC of E3-AP in 350 mM NaCl (Fig. 2b). Similarly, the PDC-E1pβ in the E3-AP in 0 mM NaCl (Fig. 2d) did not display the peak of the SEC of E1pβ-AP in 350 mM NaCl (Fig. 2a) at 14-15 ml elution volume. We inquired whether the small ~440 kDa entity in E1pβ-AP and the ~230 kDa entity in E3-AP observed only in the high ionic strength (350 mM NaCl) were forming a complex. To answer that, we subjected the peak sub-megadalton fraction of the SEC of each purification performed in 350 mM NaCl condition (Fig. 2a-b) to mass spectrometry (Fig. S2c). Surprisingly, the MudPIT analysis revealed that in E1pβ purification, E1pα and E1pβ are the primary molecules in the co-eluting entity at the elution volume of 14.5 ml. Likewise, in the E3 purification, E3 was the main molecule at the elution volume of 16 ml. Based on the MudPIT data and previous studies [14], the peak elution volume for each PDC purification at 350 mM NaCl (Fig. 2a-b) likely contained an oligomer of the bait subunit. Based on the elution volume and the MudPIT analysis, purified E1p eluted with a size of ~440 kDa and purified PDC-E3 eluted with a size of ~230 kDa.

Taken together, our data establish that the PDC is salt-labile and its integrity as a megadalton complex does not remain intact at the physiological ionic strength. Consequently, the size and the number of subunits of the complex are highly dependent on ionic strength.

### 3.3. Mammalian PDC also exhibits salt lability that is similar, but not identical, to that observed in yeast

PDC subunits E1p, E2p, and E3 in Baker yeast have conserved homology and sequence similarity with mammalian counterparts. Hence, we wanted to test whether the observed salt-lability of PDC is a unique property of yeast PDC (yPDC) or is true for the mammalian PDC (mPDC) as well. For this, HEK293T lysate was prepared and subjected to SEC in high ionic strength (350 mM). Then the size fractionated PDC components were probed by immunoblotting (Fig. 3a). As in yeast, the mammalian E1p, mE1pα (PDHA1) and mE1pβ (PDHB1), and mE3 components were detected mostly in sub-megadalton size fractions (~230 kDa) in the high ionic strength. Interestingly, the mE2p eluted only as a megadalton complex (≥ 2 MDa) even in the high salt condition. When the same size-fractions were probed for a well characterized high molecular weight SWI-SNF complex by immunoblotting (Fig. S3), its components BRG1 and BAF155 were found in megadalton size (1.3-1.5 MDa) fractions as expected. This result suggests that mPDC components are more salt-labile than the SWI-SNF complex components.

**Figure 3.**
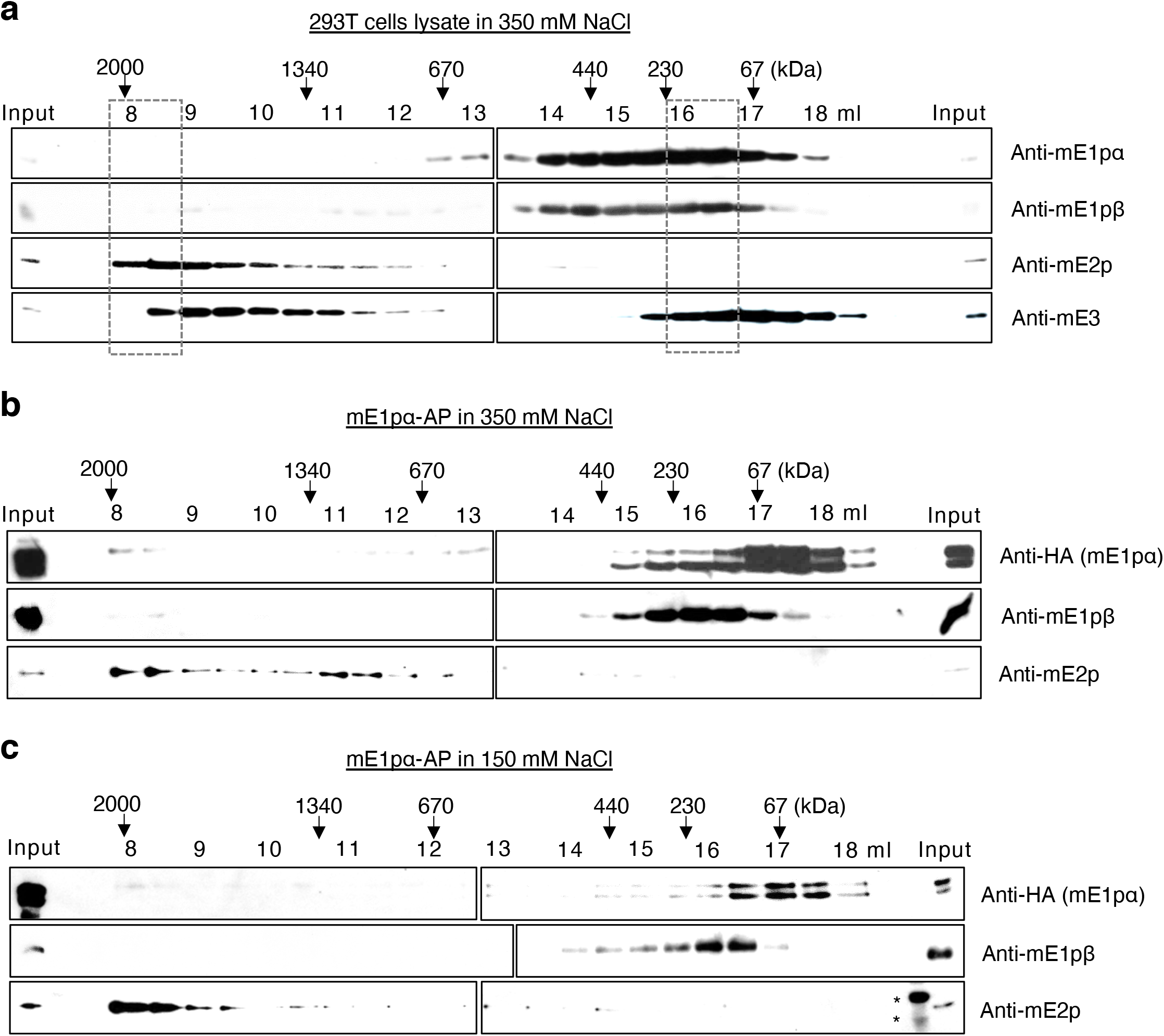
SEC of Mammalian PDC exhibits salt-lability that is similar, but not identical, to that observed in yPDC. (a) Size fraction profiles of lysates of HEK293T cells in a buffer containing 350 mM NaCl. (b) Size fraction profiles of Halo affinity purified (AP) mE1pα construct from HEK293T cells in a buffer containing 350 mM NaCl. (c) Size fraction profiles of affinity purified mE1pα construct from HEK293T cells in a buffer containing 150 mM NaCl. Asterisks denote non-specific signals. Elution volumes in the SEC via Superose 6 column for every other 500 μl fraction are indicated with the expected size of eluted proteins based on standard proteins. Boxes in dashed lines highlight the peak fractions.

To verify whether the co-fractionated mPDC components (Fig. 3a) were forming a complex, purification of mPDC via mE1pα (PDHA1) was performed from HEK293T cells. The location of the fused protein (HaloTag) for the purification was determined based on the localization of the C-terminally YFP-tagged (PDHA1-YFP) and the N-terminally GFP-tagged (GFP-PDHA1) mE1pα (Fig. S4a). The microscopy images revealed that C-terminal tagging allows the correct localization of the PDC component as suggested by the colocalization of PDHA1-YFP with a mitochondrial staining dye mitotracker, while the N-terminal tagging resulted in pan-cellular localization likely due to the unavailability of the mitochondrial localization sequence at the N-terminus of the precursor of the mE1pα [41].

Consequently, the mPDC was purified via PDHA1-HA-Halo construct from HEK293T cells in a buffer containing 150 mM NaCl and subjected to mass spectrometry (Fig. S4b). The MudPIT analysis identified all the subunits of mPDC, suggesting the intact complex was purified. When the purification of the presumably-mislocalized Halo-PDHA1 was subjected to MudPIT analysis (Fig. S4c), the amount of co-purified PDC subunits was significantly less as expected. Next, the mPDC purified via PDHA1-HA-Halo was subjected to SEC in high ionic strength (350 mM NaCl) and the eluted fractions were probed by immunoblotting against its components (Fig. 3b). The purified mPDC essentially displayed the same profile to that acquired from the cell lysate (Fig. 3a). The purified mE1pα, mE1pβ and mE2p (Fig. 3b) were eluted at 8-9 ml and 16-17 ml elution volume which matched well with those in the SEC of cell lysate (Fig. 3a). mE1pα and mE1pβ were mostly co-eluted at fractions for a sub-megadalton (~ 230 kDa) protein, likely as mE1pα2β2 tetramers in this high ionic strength condition. With an elution peak at the void volume, the mE2p seemed to only co-elute with a complex of several megadaltons (≥ 2 MDa) in size. These results suggest that mE1p is salt-labile while mE2p core is stable in high ionic strength.

Next, we inquired how the mPDC would behave in the physiological condition at 150 mM NaCl. To address that, the mPDC was purified via PDHA1-HA-Halo from HEK293T cells in a buffer containing 150 mM NaCl and subjected to SEC. The size-fractions in 150 mM NaCl were collected and probed for the mPDC components by immunoblotting (Fig. 3c). As observed for the yPDC, the mPDC displayed a similar elution profile in this physiological condition as it did with the high ionic strength (350 mM NaCl) (Fig. 3b). The mE1pα and mE1pβ co-eluted in a sub-megadalton entity of ~230 kDa, while the mE2p eluted at 8-9 ml elution volume with a complex of several megadaltons. The mPDC exhibited a striking difference as compared to yPDC in that the mE2p was more stable as a megadalton complex in high ionic strength. However, mE1p also exhibited a sub-megadalton size in a physiological condition like yE1p. The mE3 in the mE1pα-AP was below detection level in western blots (not shown). These results suggest that the salt-lability is conserved from yeast to human and might be a universal property of E1p, while the lability of E2p may differ between organisms.

### 3.4. The catalytic activity of PDC changes with ionic strength

Thus far, we have established that the PDC is a salt-labile complex compared to other protein complexes such as the SWI/SNF which maintains its integrity in high salt conditions (Fig. S3). This finding led us to speculate whether the salt lability of the complex affects its activity. Previously, a non-canonically smaller PDC observed in the process of reassembling or disassembling canonical PDC showed lower catalytic activity proportional to its size [42,43]. Thus, we postulated that ionic strength, with which the size of the PDC can vary, can modulate the activity of the PDC. To test this hypothesis, we performed *in vitro* assay of catalytic activity of the PDC. The yPDC was affinity-purified in a low salt condition (50 mM NaCl) from a PDC kinase mutant via E1pβ (*pkp2*Δ*/*PDB1-5xFLAG) to account for the highly phosphorylated inactive PDC in yeast [44]. To examine the effect of ionic strength on the PDC activity, the assay was performed with a low (50 mM NaCl) and a high (350 mM) ionic strength.

Iodonitrotetrazolium chloride (INT)-salt adduct produced by the catalytic activity of PDC was detected by absorbance at 490 nm over a time course (See material and method for details). Indeed, a lower activity was detected in the PDC in higher ionic strength (350 mM NaCl) compare to that in lower ionic strength (50 mM NaCl) (Fig. 4a). In the control experiment in which the PDC was not added in the assay, the absorbance at 490 nm was constant in both ionic strength conditions. The amount of PDC used in the two assays was verified to be the same by immunoblotting of the E1pβ-FLAG (Fig. S5).

**Figure 4.**
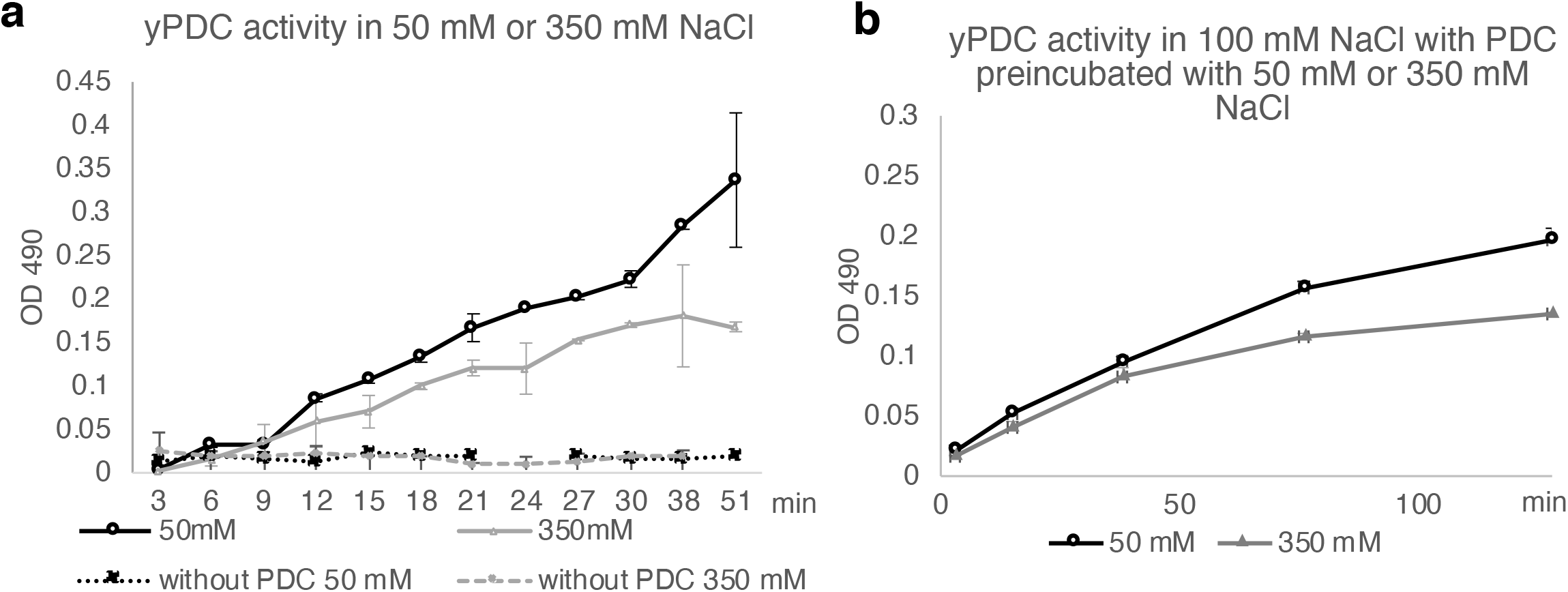
The labile character of PDC is correlated with catalytic activity. The catalytic activity of yPDC measured by absorbance at 490 nm via INT in the presence of (a) 0.6 mM INT and 50 mM or 350 mM NaCl or (b) 0.4 mM INT and 100 mM NaC with yPDC preincubated with 50 mM or 350 mM NaCl. Error bars indicate standard deviations of three replicates.

Next, to corroborate that it was the ionic strength imposed on the PDC that led to the change in the activity, we performed the assay in the same ionic strength (100 mM NaCl), while the PDC was pre-incubated in a low (50 mM NaCl) or a high (350 mM NaCl) ionic strength condition prior to the assay (Fig. 4b). Indeed, the decrease in the activity was observed when the PDC was exposed to a higher ionic strength prior to the assay. This suggested that the high ionic strength decreased the activity of PDC from that with low ionic strength. Taken together, these results suggest that the ionic strength that affects the size of PDC can modulate its catalytic activity possibly by changing the number of subunits within the complex.

## 4. Discussion

The present study demonstrates the lability of the multienzyme PDC. Several reports have shown that the PDC size, structure, and the number of its subunits can vary up to ~17 % [29,30,45–48]. However, the present study is the first report that in a systematic way establishes the lability of the PDC that forms sub-megadalton structure in physiological ionic strength.

The E1p is known to form a tetramer (E1pα2β2), which is calculated to be ~180 kDa, within the yPDC. Interestingly, the yE1p we purified eluted at fractions for ~440 kDa proteins (Fig. 2a) that mainly contained E1pα, E1pβ, and a nominal amount of E2p (Fig. S2c). PDC kinases and phosphatases have been shown to be associated with PDC by binding to E2p, but the regulatory proteins were not detected in our affinity purifications [49]. This suggests the possibility of a larger oligomer of yE1p. The E3 is known to form a homodimer, which is calculated to be ~120 kDa. In our affinity purification, yE3 eluted at fractions for ~230 kDa, which contained mostly E3 (Fig. S2c). This suggests a possible stable form of E3 as a tetramer as well as a dimer. These possible variations of oligomeric states of the PDC components further support our observation of the lability of the complex.

Forming a massive complex can provide cells a concentrated catalytic potential with high efficiency via substrate channeling and active-site coupling [14]. This notion is supported by a reassembled PDC with ~42-48 E1p heterotetramers and 6-12 E3 homodimers optimally filling 60 sites of E2p core exhibiting the maximal activity of the complex [50]. However, the lability of the PDC suggests it is possible for PDC to form structures of variable sizes. Each of the sizes may be associated with certain levels of catalytic activity. Interestingly, we showed that the catalytic activity of the PDC decreases with an increased ionic strength analogous to the activity changes observed in reassembly and disassembly studies *in vitro* [42,51]. This might be because a high ionic strength would allow fewer E1p, E2p, and/or E3 subunits in the complex leading to fewer active-site couplings, hence resulting in a less efficiency or reduced activity [15,16]. Consistent with this, PDCD patients lacking a structural component, E3BP, which facilitates the interaction between E2p and E3 within the complex, display only 10-20 % of the normal activity of the PDC [52]. The decreased activity of the PDC in such patients may be due to a reduced binding between E2p and E3, resulting into smaller complexes [22]. E1p and E3 components can bind independently and competitively to the structural core formed by the E2p oligomers with different interaction property: binding of E1p is enthalpy-driven, while that of E3 is entropy-driven [20]. The varying number of E1p and E3 components can also lead to a change in the overall activity of the complex as E1p catalyzes the rate-limiting step of the overall catalysis. In fact, various combinations of the number of E1p and E3 subunits in PDC have been reported [47]. Collectively, the size and the number of component versatility of the PDC could be a means to modulate its activity. Taken together, the versatile structure of the PDC may provide a dynamic mechanism to modulate its catalytic activity.

Recent studies reported that PDC in some cells is found in the nucleus under circumstances such as in cancer and during zygotic activation in embryonic cells, potentially as a whole ~10 MDa complex [5,6]. However, it is indecipherable how a massive protein complex with a ~450 □ size could translocate from one subcellular organelle to another even when considering the possible trafficking pathway via mitochondrial-derived vesicles (MDV) of ~100 □ in size [53]. Interestingly, a rather pliable structure that can change the size of the complex provides the means to translocate the otherwise massive protein complex. Perhaps, a smaller non-conventional sized PDC complex can translocate into the nucleus.

## FOOTNOTES

The abbreviations used are:

PDC: pyruvate dehydrogenase complex
PDCD: PDC deficiency
SEC: size exclusion chromatography
TCA: tricarboxylic acid
AP: affinity purification
IP: immunoprecipitation
CoA: Coenzyme A
MudPIT: multidimensional protein identification technology
dNSAF: distributed normalized spectral abundance factors
TPP: Thiamine pyrophosphate
INT: Iodonitrotetrazolium chloride
PES: Phenazine ethosulfate
MDV: mitochondrial-derived vesicles
OGDH: 2-oxoglutarate dehydrogenase
DLD: dihydrolipoyl dehydrogenase
TCEP: Tris(2-carboxyethyl)phosphine hydrochloride
CAM: chloroacetamide

## Data availability

The MS datasets for the human and yeast analyses may be obtained from the MassIVE repository with accessions MSV000085125 and MSV000085126, respectively (with password: JLE01146) and from the ProteomeXChange with accessions PXD018125 and PXD018126.

## Acknowledgements

We thank Workman Lab members and Stowers core facilities for their support for this work. This work was supported by funding from Stowers Institute for Medical Research and National Institutes of General Medical Sciences grant R35GM118068.

## Author contributions

Jaehyoun Lee: Conceptualization, methodology, Writing-Original draft preparation, Investigation. Seunghee Oh: Methodology, Writing-Reviewing and Editing. Saikat Bhattacharya: Methodology, Visualization, Writing-Reviewing and Editing. Ying Zhang: Methodology, Visualization, Laurence Florens: Methodology, Data curation, Writing-Reviewing and Editing. Michael P. Washburn: Methodology, Resources. Jerry L Workman: Conceptualization, Supervision, Project administration, Funding acquisition, Writing-Reviewing and Editing.

## Funding and additional information

Stowers Institute for Medical Research and National Institutes of General Medical Sciences grant R35GM118068 (J.L.W.). The content is solely the responsibility of the authors and does not necessarily represent the official views of the National Institutes of Health

