## Supplementary information for "The plasticity of the pyruvate dehydrogenase complex confers a labile structure that is associated with its catalytic activity"

**Figure S1**

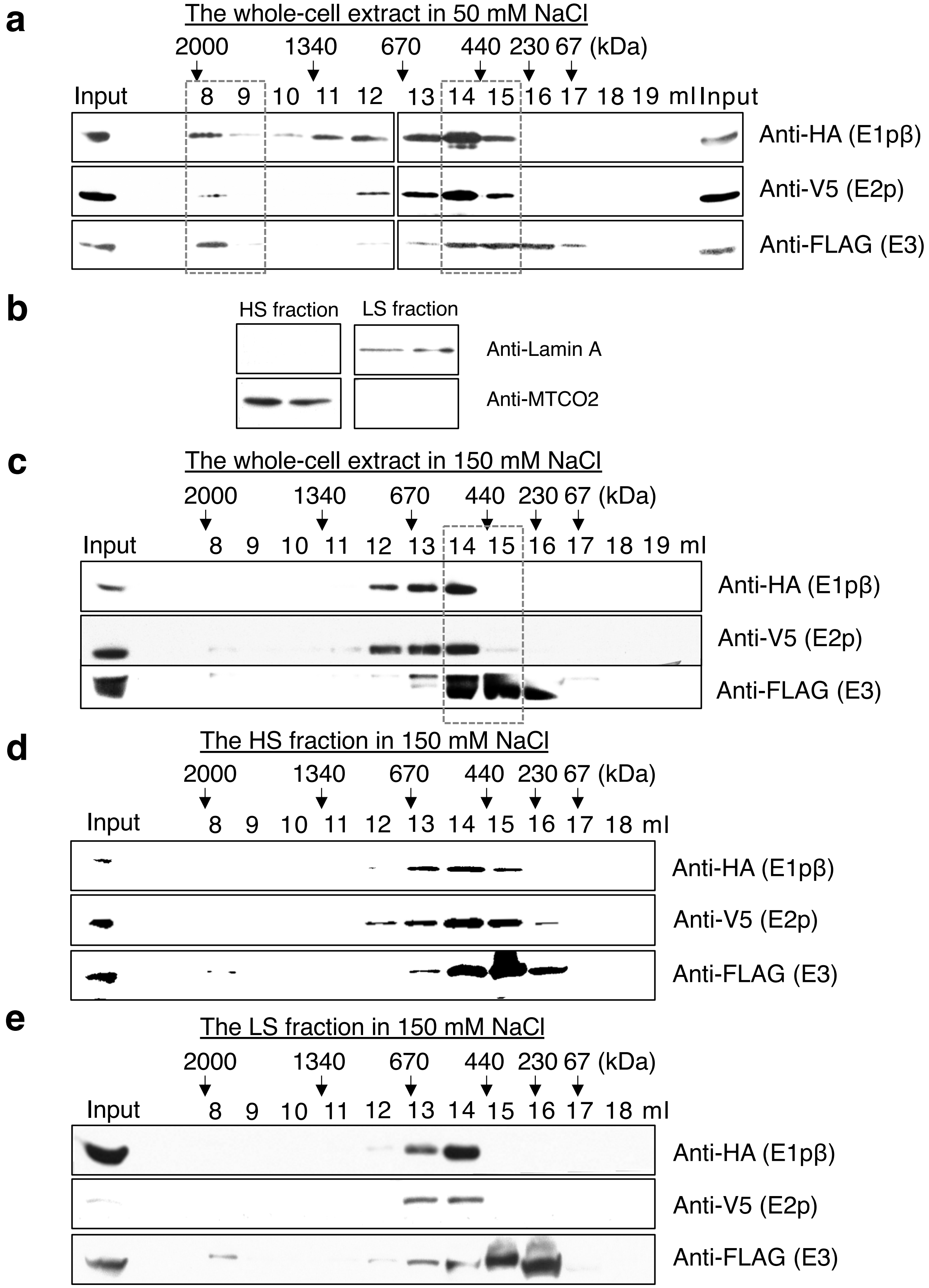


**Figure S1. The SEC profiles of yeast extract display sub-megadalton PDC components**

(a) Size fraction profiles of yPDC components of Pdb1-3xHA/Lat1-V5/Lpd1-5xFLAG strain from whole-cell extract in a buffer containing 50 mM NaCl. (b) Detection of standard proteins, Lamin A for nuclei and MTCO2 for mitochondria as a subcellular fractionation control in mitochondria-enriched HS and nuclei-enriched LS fractions from two biological replicates. For (c)-(e), Size fraction profiles of yPDC components of Pdb1-3xHA/Lat1-V5/Lpd1-5xFLAG strain from (c) whole-cell extract in a buffer containing 150 mM NaCl, (d) the HS fraction in a buffer containing 150 mM NaCl, and (e) the LS fraction in a buffer containing 150 mM NaCl. Elution volumes in the SEC via Superose 6 column for every other 500 µl fraction are indicated with the expected size of eluted proteins based on standard proteins. Boxes in dashed lines highlight the peak fractions.

**Figure S2**

**a**

| Detected Protein \ yPDC-AP bait | dNSAF E1pβ-FLAG AP | dNSAF E3-FLAG AP |
|---|---|---|
| E1pβ (Pdb1) | 0.3258 | 0.009001 |
| E1pα (Pda1) | 0.2092 | 0.006754 |
| E2p (Lat1) | 0.02917 | 0.01263 |
| E3 (Lpd1) | 0.1130 | 0.3709 |
| E3BP (Pdx1) | 0.006775 | 0.02846 |

**b**

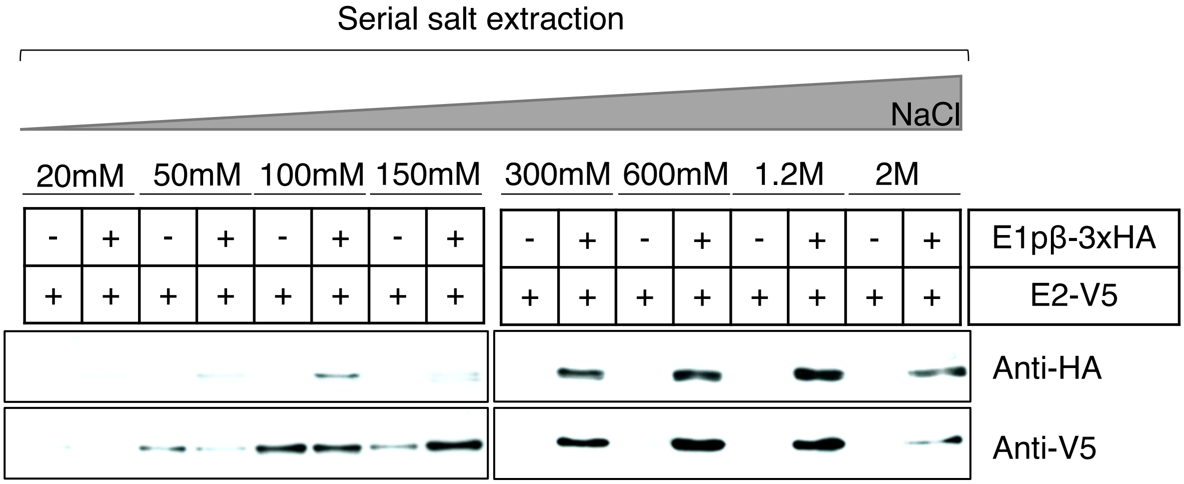


**c**

| Detected Protein \ peak fraction for each AP | dNSAF yE1pβ-AP SEC @ 14.5 ml | dNSAF yE3-AP SEC @ 16 ml |
|---|---|---|
| yE1pβ (Pdb1) | 0.3458 | 0.006628 |
| yE1pα (Pda1) | 0.3177 | 0.005217 |
| yE2p (Lat1) | 0.02249 | 0.0007306 |
| yE3 (Lpd1) | 0.0008900 | 0.6789 |
| yE3BP (Pdx1) | 0.0009750 | 0.004581 |

**Figure S2. Affinity purification of yPDC subunits suggests the lability of the complex**
(a) Relative protein levels (dNSAF) of yPDC components found in MudPIT analyses of the FLAG purified yPDC via its subunit E1β (Pdb1) and E3 (Lpd1) from Pdb1-5xFLAG and Lpd1-5xFLAG, respectively. (b) Serial salt-extraction of yPDC bound on anti-HA beads via Pdb1-3xHA in EM buffer (10 mM MOPS pH7.2, 1mM EDTA, 1 % Triton-X100). The concentration of NaCl in each washing is designated. (c) Relative protein levels (dNSAF) of yPDC components found in MudPIT analyses of a peak fraction of SEC following E1β-AP and E3-AP in 350 mM NaCl condition at the designated elution volume.

Figure S3

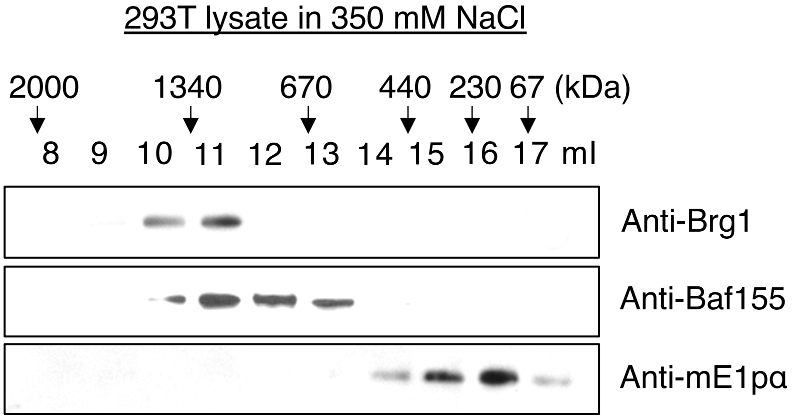


**Figure S3. SEC of HEK293T lysate suggests the salt-lability of mPDC in contrast to SWI-SNF subunits**

SEC profiles of mE1pα and mSWI-SNF subunits, BRG1 and BAF155 of HEK293T lysate in a buffer containing 350 mM NaCl. Elution volume in the SEC via Superose 6 column for every other 500 µl fraction is indicated with the expected size of eluted proteins based on standard proteins.

### Figure S4

**a**

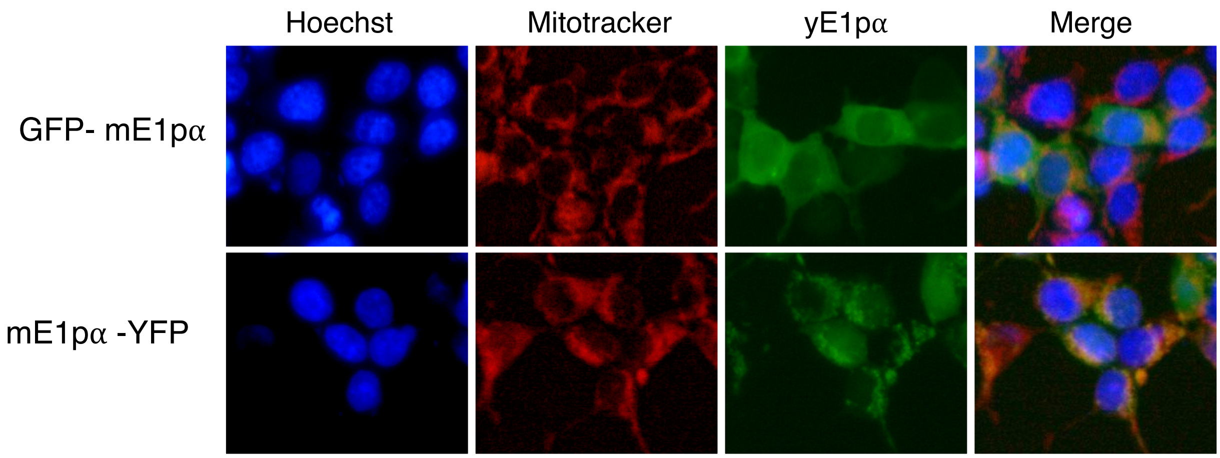


**b**

| dNSAF | mE1pα-Halo |
|---|---|
| mE1pα (PDHA1) | 0.1995 |
| mE1pβ (PDHB1) | 0.03162 |
| mE2p (DLAT) | 0.01221 |
| mE3 (DLD) | 0.001022 |
| mE3BP (PDHX) | 0.001654 |
| PDHA2 | 0.001112 |

**c**

| dNASF | Halo-mE1pα |
|---|---|
| mE1pα (PDHA1) | 0.1531 |
| mE1pβ (PDHB1) | 0.004963 |
| mE2p (DLAT) | 0.000417 |
| mE3 (DLD) | 0 |
| mE3BP (PDHX) | 0 |
| PDHA2 | 0.000164 |

**Figure S4. C-terminal but not N-terminally tagging allows correct localization of mPDC-E1α (PDHA1) and the purification of mPDC**

(a) Microscopy of HEK293T cells expressing GFP- E1α or E1α -YFP construct. For (b) and (c) relative protein levels (dNSAF) of mPDC components found in MudPIT analysis of (b) E1α-Halo and (c) Halo-E1α affinity purification.

**Figure S5**

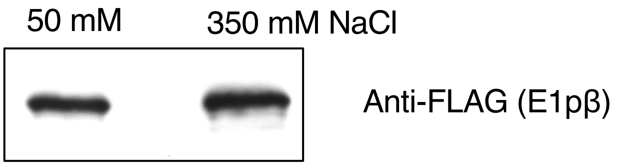


**Figure S5. Contrl experiment for catalytic activity assay**
The amount of yPDC used in the catalytic activity assay for figure 4a measured by western blot against FLAG.

**Table S1. Yeast strains used in this study**

| Name | Genotype | Source |
|---|---|---|
| Wildtype (By4741) | MATa his3Δ1 leu2Δ0 met15Δ0 ura3Δ0 pdb1Δ::KanMX4 | Open Biosystems (#YSC1048) |
| E1pβΔ (*pdb1Δ*) | MATa his3Δ1 leu2Δ0 met15Δ0 ura3Δ0 pdb1Δ::KanMX4 | Open Biosystems (#3361) |
| E2pΔ (*lat1Δ*) | MATa his3Δ1 leu2Δ0 met15Δ0 ura3Δ0 lat1Δ::KanMX4 | Open Biosystems (#7218 ) |
| E3Δ (*lpd1Δ*) | MATa his3Δ1 leu2Δ0 met15Δ0 ura3Δ0 lpd1Δ::KanMX4 | Open Biosystems (#5635) |
| E1pβ-5xFLAG (Pdb1-5xFLAG) | MATa his3Δ1 leu2Δ0 met15Δ0 ura3Δ0 Pdb1-5xFLAG::NatMX4 | This study |
| E2p-5xFLAG (Lat1-5xFLAG) | MATa his3Δ1 leu2Δ0 met15Δ0 ura3Δ0 Lat1-5xFLAG::NatMX4 | This study |
| E3-5xFLAG (Lpd1-5xFLAG) | MATa his3Δ1 leu2Δ0 met15Δ0 ura3Δ0 Lpd1-5xFLAG::NatMX4 | This study |
| E2p-V5/ E3-5xFLAG (Lat1-V5/Lpd1-5xFLAG) | MATa his3Δ1 leu2Δ0 met15Δ0 ura3Δ0 Lat1-V5::HIS3 LPD1-5xFLAG::NatMX4 | This study |
| E1pβ-3xHA/ E2p-V5/ E3-5xFLAG (Pdb1-3xHA/Lat1-V5/ Lpd1-5xFLAG) | MATa his3Δ1 leu2Δ0 met15Δ0 ura3Δ0 Pdb1-3xHA:KanMX4 Lat1-V5:His3 Lpd1-5xFLAG:NatMX4 | This study |
| *pkp2Δ*/PDB1-5xFLAG | MATa his3Δ1 leu2Δ0 met15Δ0 ura3Δ0 pkp2Δ::KanMX4 Pdb1-5XFLAG::NatMX4 | This study |
